# Cell substrate patterns driven by curvature-sensitive actin polymerization: waves and podosomes

**DOI:** 10.1101/2020.02.17.953018

**Authors:** Moshe Naoz, Nir S. Gov

**Affiliations:** Weizmann Institute of Science

**Keywords:** Actin waves, curved proteins, dynamic instability, podosomes

## Abstract

Cells adhered to an external solid substrate are observed to exhibit rich dynamics of actin structures on the basal membrane, which are distinct from those observed on the dorsal (free) membrane. Here we explore the dynamics of curved membrane proteins, or protein complexes, that recruit actin polymerization when the membrane is confined by the solid substrate. Such curved proteins can induce the spontaneous formation of membrane protrusions on the dorsal side of cells. However, on the basal side of the cells, such protrusions can only extend as far as the solid substrate and this constraint can convert such protrusions into propagating wave-like structures. We also demonstrate that adhesion molecules can stabilize localized protrusions, that resemble some features of podosomes. This coupling of curvature and actin forces may underlie the differences in the observed actin-membrane dynamics between the basal and dorsal sides of adhered cells.

## 1. Introduction

The actin cortex of cells is the prominent driver of membrane shape deformations, which exhibit a huge variability, from propagating waves to stable protrusions. It is often observed that the actin-membrane dynamics of adhered cells is very different between the basal and dorsal sides. The main difference between the two side is that on the basal side the membrane is held at close proximity to the solid substrate, while on the dorsal side the membrane is usually free to deform into the surrounding fluid. In this paper we explore theoretically the actin-membrane dynamics in the presence of the confinement of the substrate, when the actin polymerization is nucleated by curved membrane complexes.

Cells exhibit a variety of propagating waves of actin polymerization on their basal plasma membrane, which are observed under many conditions such as the initial formation of adhesion [1] and during cell motility [2,3,4]. When these waves propagate on the dorsal side of an adhered cell, or along its perimeter edge, they are accompanied by large membrane deformations. However, when these waves propagate along the basal membrane, at the interface between the cell and the underlying solid substrate, such membrane deformations have not been unambiguously observed. These basal actin waves have been studied intensively [5,6,7,8], and many of their features exposed. Mostly these waves have been treated in the framework of reaction-diffusion models [8], where membrane deformations do not play a role.

In previous works [9,10] we have investigated theoretically and experimentally the possible role of curved activators of actin polymerization in the propagation of membrane-actin waves. In these works the positive feedback is in the form of an actin nucleator that has a convex shape (such as the I-BAR protein IRsp53 for example [11,12]), such that it tends to accumulate at the tips of membrane protrusions that are driven by the actin polymerization force. The negative feedback, which is necessary for wave propagation, can be provided by the contractile force of myosin-II [9] or the recruitment of concave-shaped actin nucleators (such as the BAR family proteins, Tuba for example [13]) [10]. More recent work proposed that the negative feedback for propagating basal actin waves arises from the actin network itself [14].

In this paper we explore the dynamics of the membrane-actin system in the presence of only the convex nucleator, but in the presence of a confining boundary which represents the effect of the solid substrate. When there is no confinement, our model predicts that this system can become unstable and drive the spontaneous initiation of membrane protrusions through a Turing-type instability [15,16,17,18,19], as is indeed observed in experiments [20,21,22,23]. We show that in the presence of a confining boundary this system indeed supports protrusions, which are however modified compared to those growing on a free membrane: protrusions may split, and may even convert into propagating rings. These theoretical results may explain some puzzling features observed for actin waves that propagate at the substrate-attached cell surface, such as their tendency to form doublets of concentric actin fronts [8,24,25].

In the last section we demonstrate that by adding adhesion of the membrane to the substrate localized protrusive structures can be stabilized, and these share some features with localized adhesion structures called podosomes.

## 2. Results

### 2.1. Expanding ring of membrane-actin wave

Our model is based on the description of the membrane shape in terms of a single height variable *h*(*x, y*), which is appropriate for small membrane deformations (Monge gauge). This is applicable for the present system, where the membrane is adhered to a solid substrate that confines the extent of its normal deflection. On the membrane we consider a density field *n*(*x, y*) of “activator” proteins, which are both curved and recruit the polymerization of actin. These “proteins” may therefore represent bound complexes that contain several proteins, that together have this combination of properties. The curved membrane complexes can diffuse on the fluid membrane, as well as form high density aggregations.

We solve the equations of motion for the two fields (10,11) numerically using an explicit finite difference scheme with periodic boundary conditions. We first investigate the response of the system to a single small gaussian perturbation (see Supplementary Movie S1, Fig.1). We choose values for the parameters of the model such that we are in the unstable regime, and protrusions grow spontaneously. However, the numerical values of these parameters are not fitted to any particular experimental measurement, and are simply chosen to demonstrate the qualitative behavior over realistic length and time scales.

**Figure 1.**
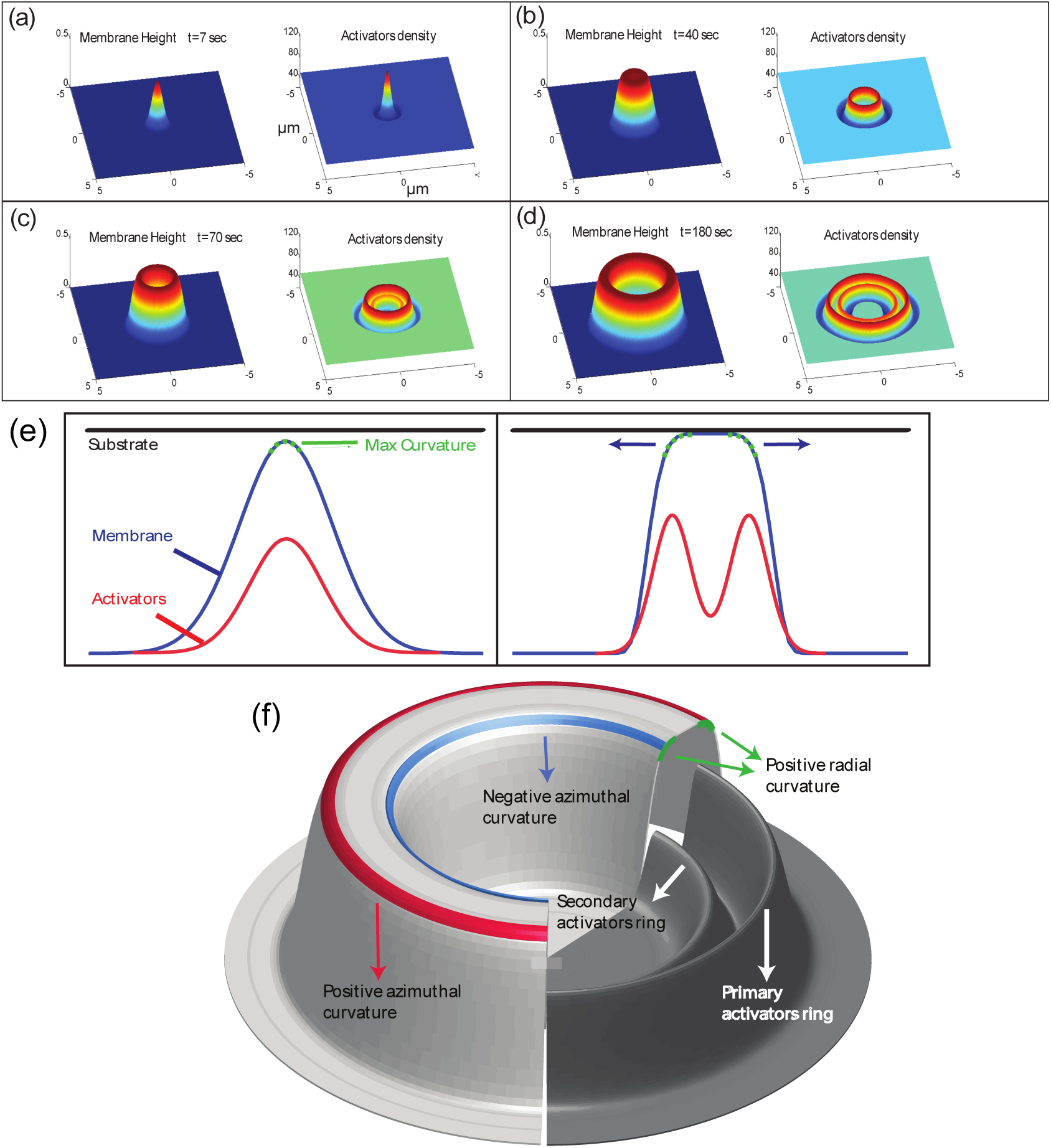
(a-d) Numerical integration of equations (10,11) over a period of 3 minutes, over a membrane segment of size 10 × 10*µ*m^2^. The parameter values used: *D* = 0.1*µ*m^2^/s, 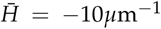,n_0_ = 50*µ*m^−2^, *n*_*s*_ = 300*µ*m^−2^, *A* =3.8·10^−5^ *kgµ*m^5^sec^−2^, *κ* = 20k_*B*_T, *s* =8.28·10^−5^ *kgµ*m^4^sec^−2^, *µ* =1.66·10^6^sec*µ*m^−2^kg^−1^, *h*_wall_ = 0.5*µ*m. (a) Initial growth of the protrusion, prior to contact with the substrate. Note that throughout the paper the plots of the “activator density” is with respect to the background, uniform density *n*-*n*_0_. (b) The protrusion after it comes into contact with the substrate and the membrane at the tip becomes flat. As a consequence an activator ring are formed at the edge of the membrane protrusion, where there is large positive curvature. (c) The membrane at the disk center has retracted back towards the unperturbed position at *h* = 0, and a secondary inner ring of activators begins to form. (d) The membrane ring and two activator rings expand further. (e) An illustration of the mechanism that drives the expansion of the protrusion radially outwards. When the membrane reaches the flat substrate its curvature diminishes and the activators are then aggregated at the location of the highest curvature - the shoulders. However, since once the activators aggregate they push the membrane against the substrate which results in the flattening of the shoulders and the formation of new shoulders further away from the protrusion center. (f) A diagram of the structure of the expanding ring. Marked in green are the regions high in curvature in the radial direction which is similar in magnitude for both the inner and the outer rings. Marked in red is the outer radius curvature in the azimuthal direction which is positive and marked in blue is the inner radius azimuthal curvature which is negative. Also shown is the concentration of activators which aggregate into two rings at the outer and inner radii of the membrane ring. The concentration of the outer activators ring is higher than the concentration at the inner ring due to the different azimuthal curvatures.

As shown in Fig.1a, the perturbation initially grows into a protrusion with a lateral width of the order of the most unstable wavelength *λ*_*c*_ (Eq.12). During this growth stage, the protrusive force of the actin locally squeezes the layer of long molecules (glycocalix) that buffers the outer surface of the membrane from the substrate (Fig.8). Due to the positive feedback, the density of activators increases at the tip of the growing protrusion (note that throughout the paper the plots of the “activator density” is with respect to the background, uniform density *n*-*n*_0_).

**Figure 2.**
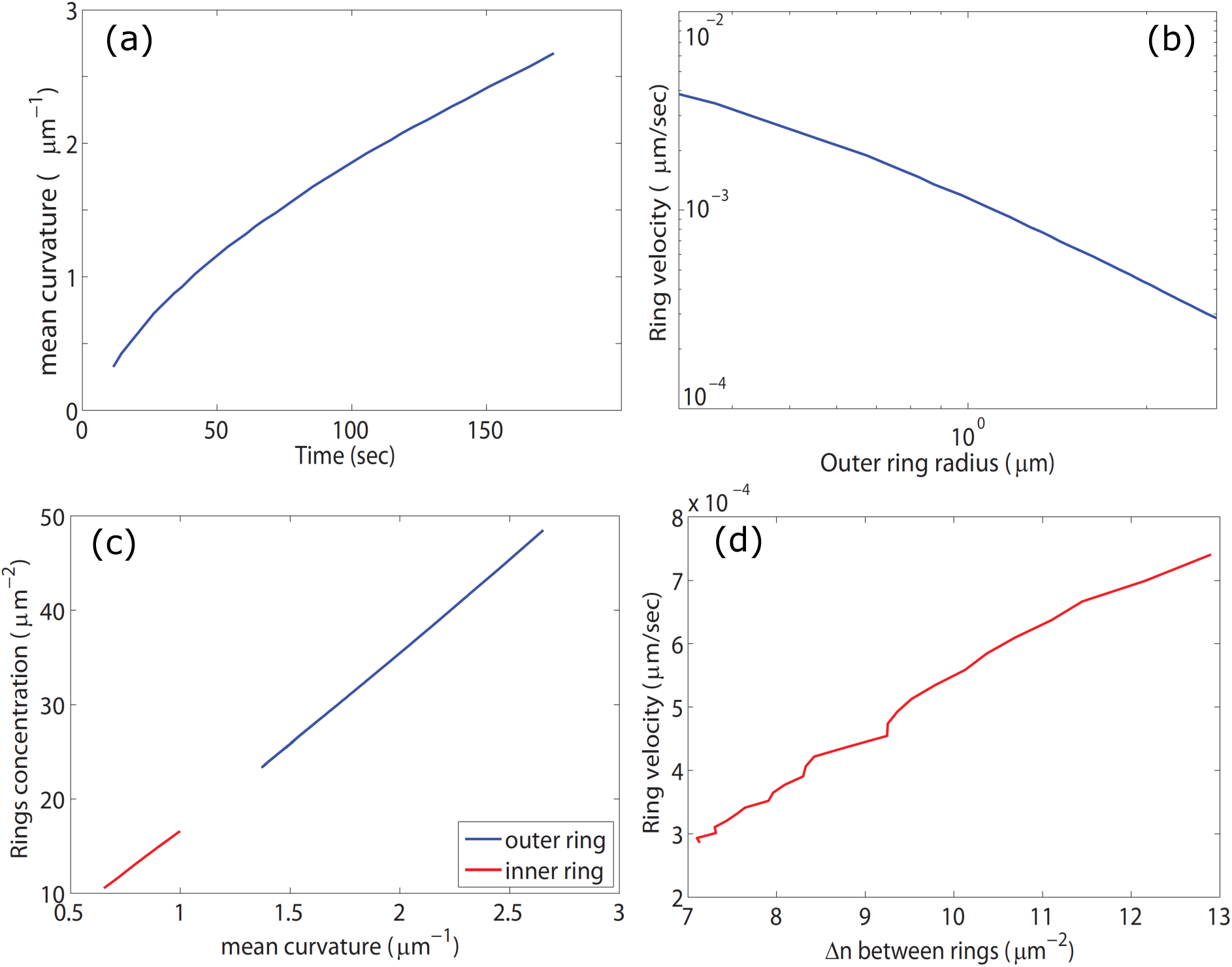
(a) The radius of the outer membrane disk (and later ring) as a function of time (all panels correspond to the simulation shown in Fig.1). (b) A log plot of the expansion speed of the outer radius vs the radius. We see that the graph is curved at small radii (where the ring is actually a disk) but as the radius grows (and the ring forms) the graph approaches a straight line indicating the power law relation: *V*_ring_ ∼ 1/*R*. (c) The peak values of the differential concentration *n*-*n*_0_ at the inner (red) and outer (blue) rings as a function of the local membrane curvature. We see the the concentrations are linear in the curvature, as given by Eq.3. (d) The ring outwards velocity as a function of the difference in concentrations between the inner and outer activators ring.

**Figure 3.**
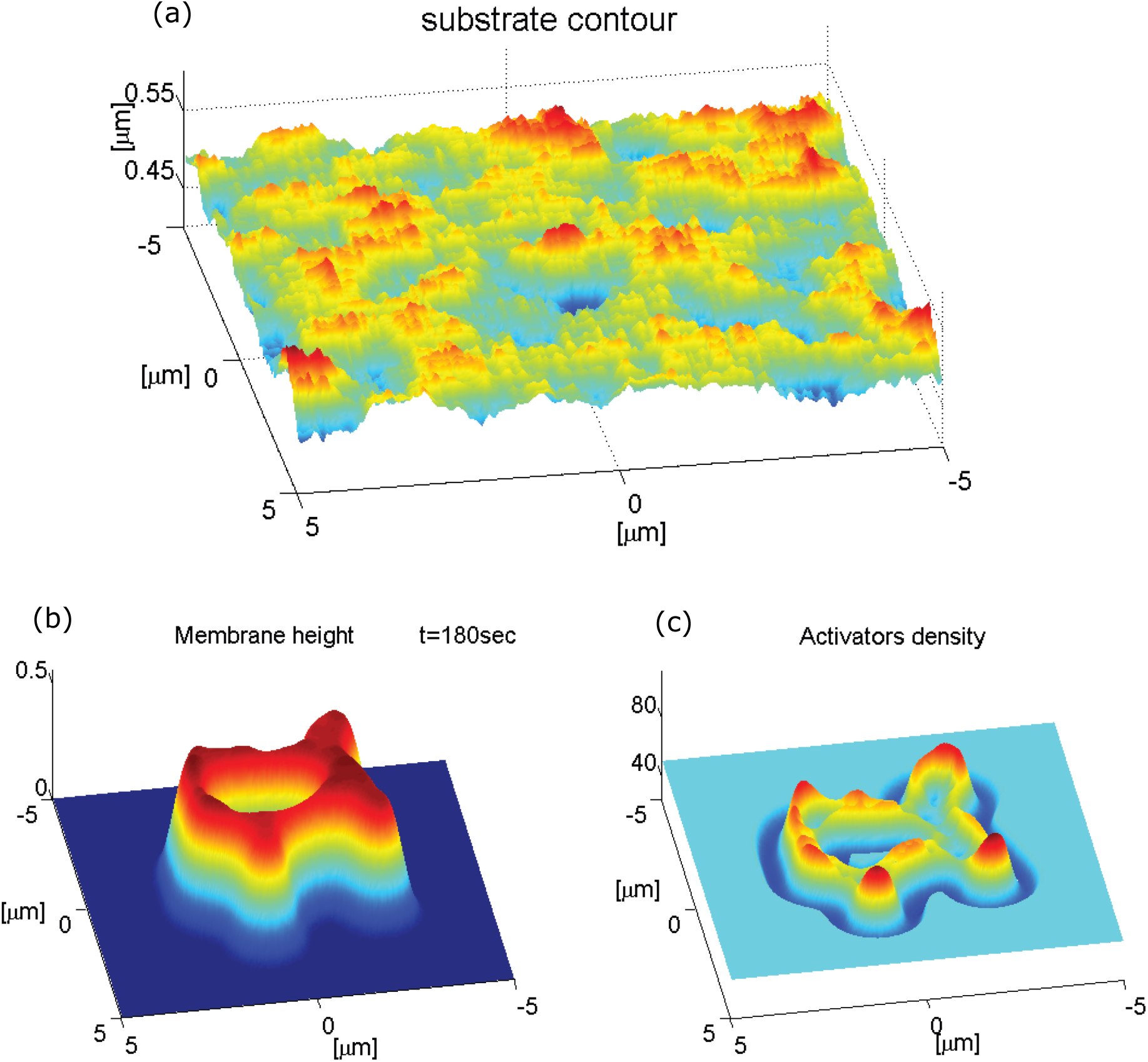
(a) Substrate with random roughness, of Gaussian amplitude, with an average amplitude of a few tens of nanometers. (b,c) Numerical integration over a period of 3 minutes of a ring expanding over a membrane segment of size 10×10*µ*m^2^. We used the same parameter values as in Fig.1.

**Figure 4.**
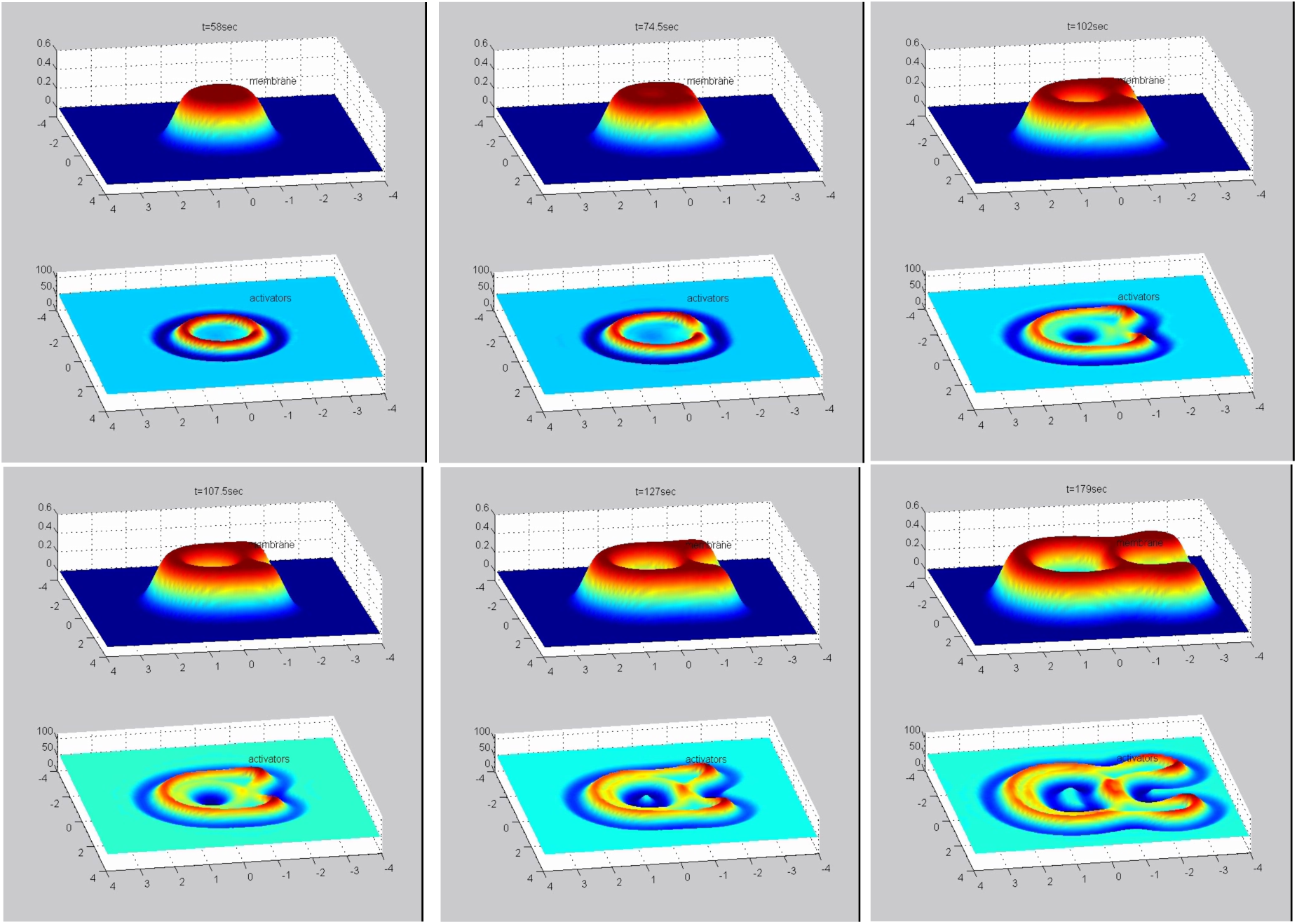
Numerical integration of a ring expanding over a membrane that contains a cylindrical ridge along the *x*-axis. From top-left to bottom-right, each panel shows the system at increasing time, with the membrane shape and activator density in the top and bottom parts respectively. We used the same parameter values as in Fig.1.

**Figure 5.**
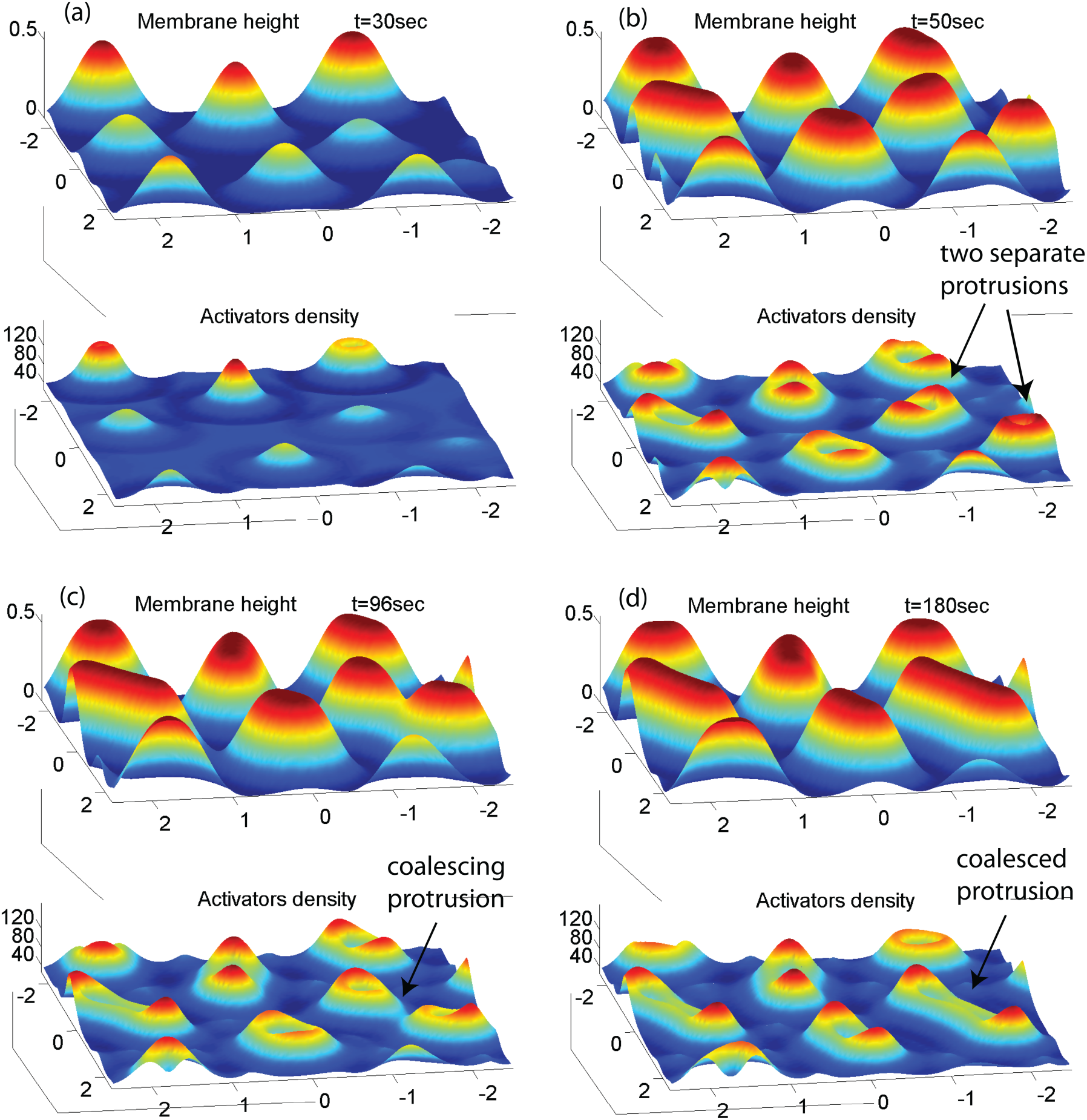
Simulation over a period of 3 minutes over a membrane segment of size 5×5*µ*m^2^, with Gaussian noise in the initial membrane height with variance of 10nm. The membrane’s random initial deformations develop into protrusions of lateral size ∼ *λ*_*c*_. (a) The initial growth period before most of them make contact with the substrate. (b)-(c) The stabilization period of the protrusions, which elongate into the available space and may coalesce with very proximal protrusions. (d) The final steady state of the system. We used the same parameter values as in Fig.1.

**Figure 6.**
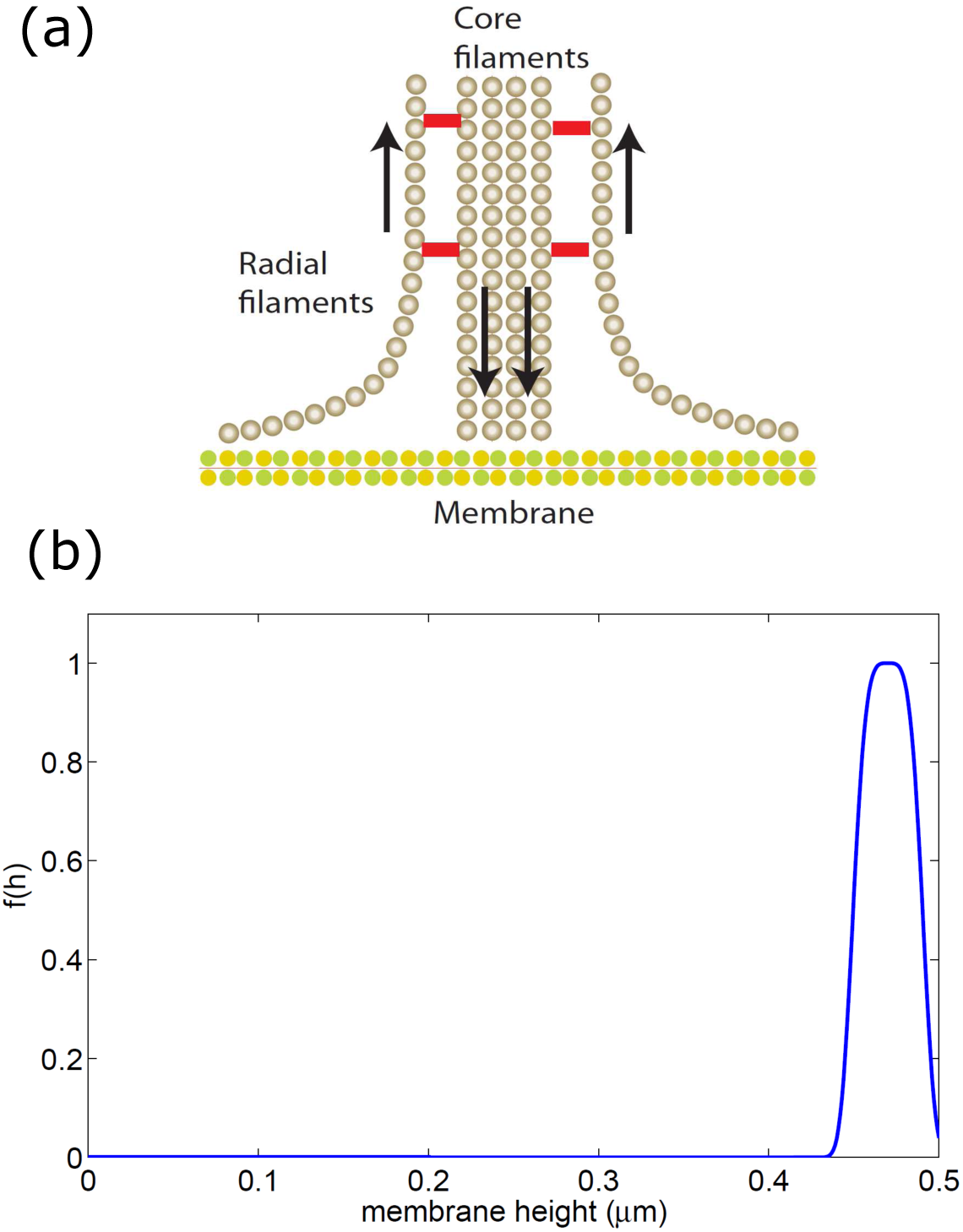
(a) Schematic illustration of the actin cables around the core of the podosome, where the treadmilling flow induces shearing and pulling forces near the membrane. These forces are thought to activate integrin-based cell adhesion [28]. (b) The function *f*(*h*) (Eq.4) that we used to limit the binding of the adhesion proteins only to within a distance of 2*l*_0_ from the substrate (here *h*_wall_ = 0.5*µ*m).

**Figure 7.**
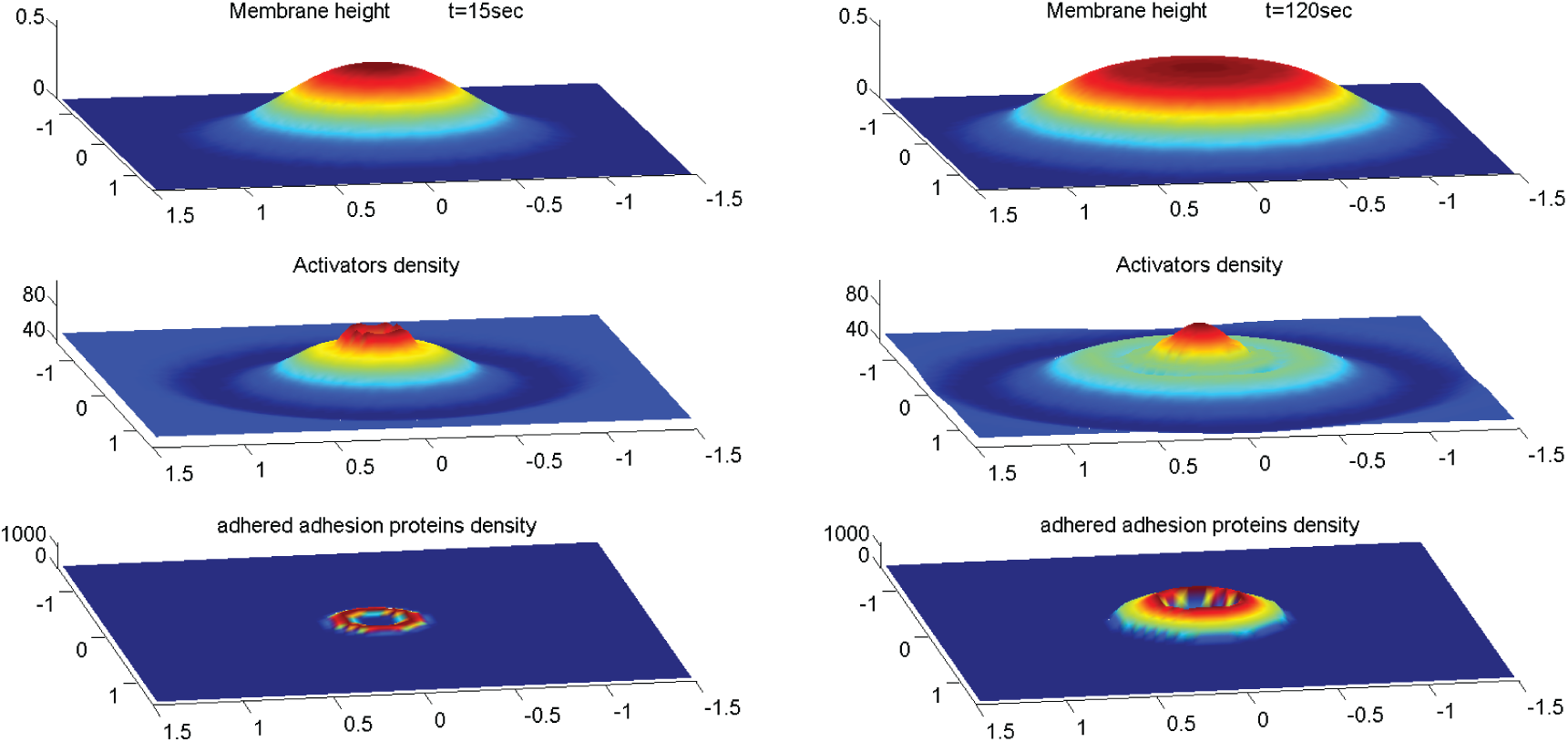
Numerical integration of equations (15) for the case of constitutive active activators over a period of 2 minutes over a membrane segment of size 5×5 *µm*^2^. The parameter values used were the same as used previously, with the following changes: 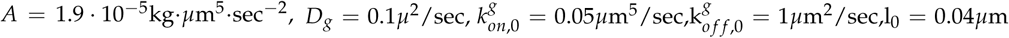 and *g*_0_ = 50*µ*m^−2^ (the initial uniform concentration of the free adhesion proteins). The perturbation develops into a protrusion and when it reaches the wall the activators at the core start expanding into a ring but this expansion is inhibited by the adhesion ring that forms around it. The actin core is then trapped by the adhesion ring and the structure stabilizes.

**Figure 8.**
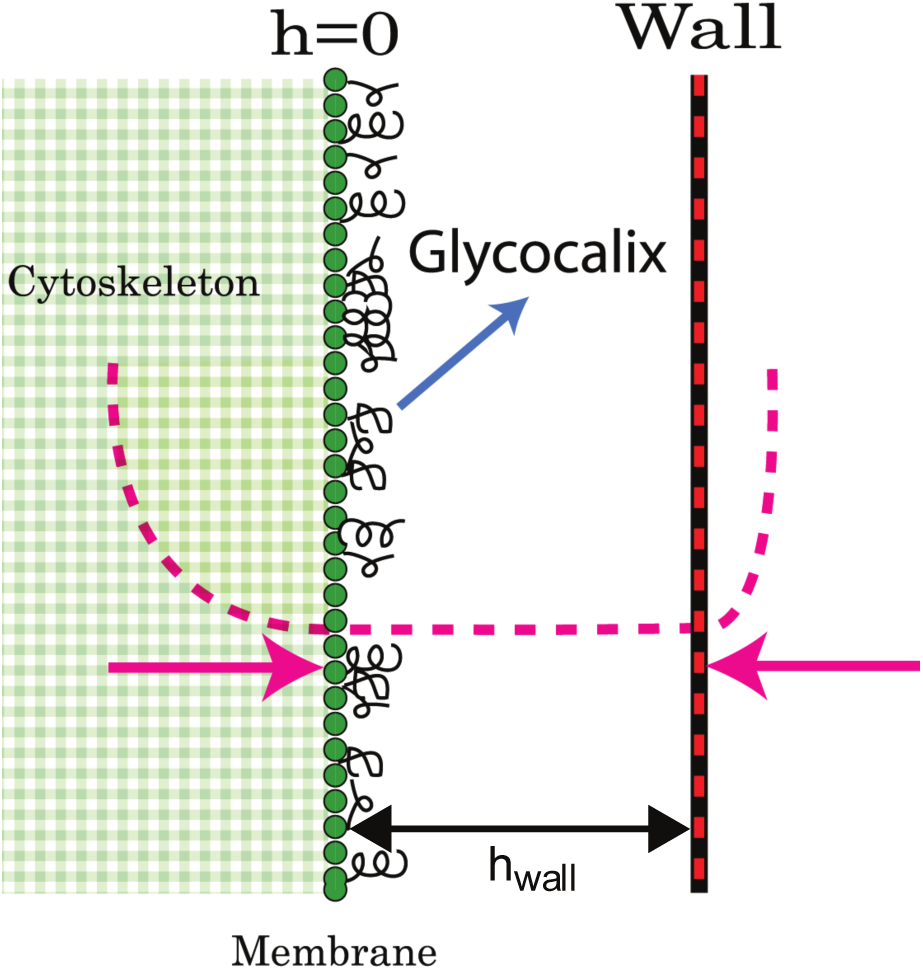
Illustration of the model. The substrate (“wall”) acts as a spring with a force *F*_wall_(*h* − −*h*_wall_) (right pink arrow) when the membrane height *h* exceeds the boundary location *h*_wall_. The cytoskeleton acts in the same way only in the opposite direction (left pink arrow) and and is typically softer, i.e. *F*_cyt_ < *F*_wall_.

Once the protrusion’s height exceeds *h*_wall_, the substrate limits further growth and the membrane shape tends to match the contour of the substrate. If the substrate is sufficiently flat the activators which were aggregated at the tip of the protrusion disperse. Note that we do not consider at this stage any adhesion to the substrate, so that the activators remain mobile on the membrane even when it is in contact with the substrate. The result is a rolling instability where the activators continually aggregate at shoulders of the protrusion (Fig.1b,e), which are the location of the highest convex curvature. The aggregation of activators increases the protrusive force exerted on the shoulders which are therefore results in the membrane deformation moving radially outwards. The rolling instability results in the protrusion developing into an expanding circular structure.

We emphasize that this model includes only normal deformations of the membrane, so the actin force does not directly push the membrane sideways along the substrate. The movement of the protrusion laterally is driven by the flow of the curved activators, and the coupling to the protrusive force of the actin polymerization. Note that a similar behavior is expected for curved activators that adsorb in a curvature-dependent manner from the cytoplasm [10].

The membrane’s initial shape is an expanding cylinder and the activators form a ring at the membrane perimeter of the protrusion (Fig.1b). Once the radius of the membrane cylinder is sufficiently large, the membrane at the center of the cylinder, which is no longer supported by a surplus of actin protrusive force, falls back to the initial height (at *h* = 0). This happens due to the inherent repulsion between the membrane and the substrate, cause by the “cushion” layer of long molecules (glycocalix) that cover the outer surface of the membrane (Fig.8). When the membrane cylinder changes into a ring shape, a secondary inner ring of activators forms at the inner shoulder of the membrane ring (Fig.1c,d,f), where there is high convex curvature.

The amplitude of the activators density at the inner ring is initially considerably smaller then the outer ring amplitude. The reason for the amplitude difference is the difference in the mean curvature between the outer an inner rings. While the radial curvatures (the curvature along the radial coordinate centered at the protrusion center) are very similar, the azimuthal curvature (the curvature along the angular coordinate), which is of the order of 1/*R* (*R* is the radius of the ring) is positive at the outer radius and negative at the inner radius (Fig.1f). Therefore at small radii, the difference in the mean curvatures is significant. The higher convex curvature at the outer ring means that the convex activators aggregate there more and the resulting protrusive force exerted on the membrane is stronger. This imbalance results in the continued outwards expansion of the ring. Note that the inner actin ring does *not* move inwards, since the curved actin activators flow towards increasing mean convex curvature, which decreases if the ring would shift to a smaller radius. Therefore the inner ring is also propagating outwards, at a velocity which is very similar to that of the outer ring, maintaining a roughly constant distance between them.

However, as the ring radius grows larger, the difference between the azimuthal curvatures at the inner and outer rings diminishes and the difference in the amplitudes of the activators rings (and therefore the protrusive force) decreases, which reduces the speed of the ring expansion (Fig.2a,b). In Fig.2c we plot the activators density at the inner and outer rings as function of the local mean curvature, and in Fig.2d we plot the ring velocity as a function of the difference between the density of activators at the inner and outer rings. The plot shows that the velocity is proportional to that difference, i.e *V*_ring_ ∝ Δ*n* = *n*_outer_ − *n*_inner_. The results indicate that the ring velocity is indeed proportional to the imbalance in the pushing force of the two actin rings, and explains why it decreases as: *V*_ring_ ∼ 1/*R* (Fig.2b).

We can quantify these observations by the following calculation: If we hold the membrane shape constant, we can solve the steady state concentration profile of the curved activators that corresponds to that shape. By taking 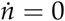 in Eq.11 and integrating once we get

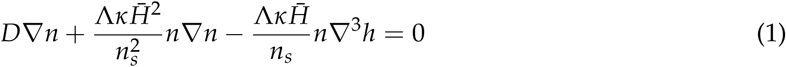

we then divide by *n* and integrate again and we are left with

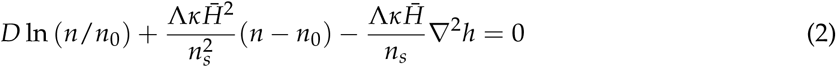

For large concentrations we can neglect the first term on the left hand side and get

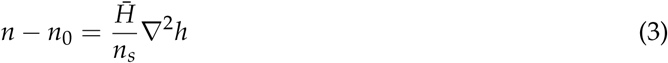

We therefore find that when the dynamics of the activators is faster than the expansion rate of the membrane deformation, so that the activators’ concentration is in a quasi steady state, we get that the activators amplitude is approximately proportional to the local membrane curvature. In Fig.2c we plot the concentration of the inner and outer rings vs. the mean curvature at these locations. The plot shows the concentrations are a linear function of the mean curvature which confirms that for the parameters used in the calculation, the dynamics of the activators is indeed faster than the membrane dynamics, and the result of Eq.3 is valid.

Since in our model the concentration of actin activators is strongly affected by the local membrane curvature, the dynamics of the ring is sensitive to the topography of the substrate. We illustrate this by simulating the dynamics on a substrate that is roughened with random bumps with an average amplitude of a few tens of nanometers (Fig.3a). Using the same set of equations and the same initial conditions as shown in Fig.1a-d, we now get the result shown in Fig.3b,c. The overall qualitative behavior is similar to the case with a smooth surface, that is, the formation of an expanding ringlike membrane structure with inner and outer rings of activators. However, the shape of the expanding structure is strongly affected by the substrate roughness and did not retain the circular symmetry it started with. The membrane ring also shows short “finger-like” protrusions extruding radially from the perimeter, which are accompanied by very strong aggregation of activators. The inner activators ring does not extend into these deformation. Due to the surface roughness, and the consequent fluctuations in the membrane curvature, the distribution of the activators inside the membrane ring-like structure becomes very inhomogeneous and fragmented.

Another illustration of the effects of substrate topography is shown in Fig.4 (Supplementary Movie S2), where a single elongated cylindrical ridge protrudes from the otherwise flat surface (along the *x*-axis). As can be seen, when the membrane ring first reaches the tip of the ridge it is curved backwards with respect to its expansion direction (top middle panel). As the membrane wraps around the protruding ridge, this membrane part develops negative curvature and the ring of actin activators breaks up at that point (top right panel). Only when the inner activators ring forms, this tip of the bump becomes favorable and concentrates activators that begin to push the membrane ring backwards towards the point of initiation (bottom right panel). On the two side of the elevated ridge the original membrane ring propagates faster than on the flat substrate. This is due to the higher concentration of actin activators, which is caused by the high positive curvature in the sharp corners that the elevated ridge makes with the flat substrate (bottom middle and right panels).

The simulations in Figs.3,4, demonstrate that the distribution of actin activators (and therefore actin filaments) can become highly fragmented due to surface undulations, while the enveloping membrane structure remains continuous and smooth. It is therefore not straightforward to relate the actin signal to the membrane topography when interpreting experimental images of such cell-substrate waves. It also shows that topographic features can cause local direction reversals and break-up of these actin waves.

### 2.2. Array of localized protrusions

We now study the conditions that may stabilize localized structures driven by the same curved activator we used so far. In Fig.5 we show that starting with different initial conditions may lead to a quasi-stable array of localized structures. A random noise in the initial membrane height or activators density, instead of a single perturbation, gives rise to multiple protrusions, each with a lateral size comparable to *λ*_*c*_ (Fig.5). When these protrusions reach the substrate they undergo the same process of flattening and expanding that we saw before for a single protrusion. During their expansion, the distance between these protrusion naturally decreases. The interaction between these protrusion is repulsive due to the positive membrane curvature region trapped between neighboring protrusions, similar to those observed in other curved membrane-bound aggregates [26]. Therefore once they have expanded and reached a distance of order *λ*_*c*_ from each other their expansion is halted and they stabilize. Over the course of the stabilization, the protrusions expand into all the space that is available by the repulsive interactions between neighbors, leading to the elongation of some of the protrusions. Furthermore, if two protrusions were initially formed in close proximity, the energy barrier between them is decreased and they can coalesce into a single elongated protrusion (arrows in Fig.5b,c,d point to such a process).

Note that in these localized protrusions the actin forms small rings within each protrusion, due to the same process we found in the expanding ring (Fig.1), on a smaller scale. These protrusions, which may seem stable, are highly sensitive to small perturbations, since they are stabilized only by their mutual repulsion due to the local membrane-driven barrier. Over long times we expect noise to cause them to shift and coalesce. Non-linear terms drive coalescence over such barriers, over long time [26].

### 2.3. Adhesion-stabilized localized protrusion: the podosome

In order to stabilize an isolated protrusion on the basal side, using the curvature-actin mechanism that we propose, we need to stabilize the localized actin core and prevent the tendency of the protrusion to expand outwards as a ring (Fig.1). An example of a localized adhesion structure on the basal size of many cell types is the podosome [27]. Podosomes are actin-rich protrusive adhesion structures formed on the membrane of several cell types, and have been implicated in the processes of cell migration, tissue invasion and extracellular matrix (ECM) degradation. The mechanisms that give rise to podosome formation, and their large-scale organization in the cell, are still poorly understood and are the subject of ongoing current research [28]. Podosomes are typically formed in monocytic cells such as macrophages, osteoclasts and dendritic cells and similar structures called invadopodia have been observed in carcinoma cells [29,30,31,32,33]. They are relatively dynamic, formed and destroyed in the span of a few minutes and are formed only on the interface between the cell and a substrate.

The podosome’s actin core is surrounded by an adhesion ring, which we did not include so far in the theoretical model, and we therefore suggest that this component may stabilize the core and prevent its ring-like expansion. We propose that the adhesion molecules form a diffusion-barrier that greatly inhibits the diffusion of the membrane-bound actin nucleators (Eq.14), and thereby stabilizes the localized core. We incorporate the adhesion into the model with the same approach that we used to incorporate the actin, namely, all the proteins involved in the adhesion process, from plaque proteins that form a scaffold around the actin core to the integrins which connect between the cellular membrane and external ligands, are grouped into a single component which we denote the “adhesion proteins”. We consider adhesion proteins to be membrane proteins that have two possible states: a non-adhered, freely diffusing state with concentration *g* _*f*_ and an adhered, immobile state with concentration *g*_*b*_. The transition from non-adhered to adhered state is only possible when the distance between the membrane and the substrate is small enough, for the membrane-bound integrins to bind to the substrate. Note that we do not explicitly describe the inter-podosome actin network [32], but rather focus on modeling a single, isolated podosome.

In addition to the geometric constraint, the binding rate of the membrane-bound adhesion proteins depends on the application of tensile forces, as integrins are known to exhibit catch-bond properties [34,35]. In the vicinity of the actin core of the podosome, the tensile force is thought to arise at the outer edge of the core, where the actin filaments flow towards the actin core and apply a pulling and shearing force that facilitates integrin adhesion [36,37,38] (Fig. 6a).

We combine the two properties that affect the adhesion listed above, in a very simplified way, in the following equations for the binding/unbinding rates of the adhesion proteins (used in the first order kinetics Eqs.15)

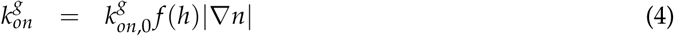

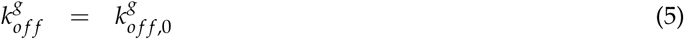

where *f* (*h*) has the profile shown in Fig.6b so that binding is only permitted close to the substrate. The last term in Eq.4 describes in the simplest way the fact that the adhesion is dependent on the spatial gradients in the actin force that is applied on the membrane. In addition, we note that the adhesion strength that determines 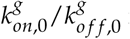 is affected by the substrate stiffness and chemical composition.

As before (Fig.1), we investigate the system’s behavior due to a small gaussian perturbation in the membrane height (Fig. 7). Numerical integration of Eqs.15 shows that the perturbation grows into a protrusion and the curved activators aggregate at the tip of the protrusion. When the protrusion reaches the substrate, the activators disperse from the center towards the shoulders. However, at the same stage, adhesion proteins bind to the substrate around the activators and inhibit their dispersion. If the adhesion proteins aggregate quickly enough they can trap the activators in the protrusion core, despite the negligible curvature that the membrane at the center posses due to the confinement by the flat substrate. However, the activators in the core may still form a small ring due to small changes in membrane curvature (Fig. 7). When we choose a shorter binding protein, we get a core of smaller radius, which is uniform, i.e. the activators do not form a ring shape. We therefore demonstrate that the adhesion-ring can indeed function as a diffusion-barrier that stabilizes the actin core. On long-time scales, where the membrane-bound actin nucleators may detach from the membrane, the actin core may decay and this process could limit the podosome life-time.

Overall, the stable podosome-like structure that our model produces exhibits many properties of the podosome. This serves to demonstrate that a model with very few components can give rise to spontaneous formation of membrane protrusions at the basal side of cells, which form an adhesion complex that closely resembles podosomes. It is sometimes observed that the actin core of podosomes may be slightly depleted at its center near the membrane [39], which is a feature that can also appear in our model (Fig.7).

Note that our model contains a single type of curved actin “activator”, while podosomes seem to posses a complex composition of actin filaments [40], as well as a complex and dynamic inter-podosome actin-myosin network [41]. Future modeling of these cellular structures, could involve more of these components, allowing for the simulation of their large-scale dynamics, where podosomes form macro structures comprised of many podosomes [38,42]. For such long-time simulations we will also need to include processes that limit the life-time of individual podosomes. In most cells, podosomes are organized in a cluster of uniform distribution with a characteristic distance between, undergoing processes of fusion and fission [43]. This organization and dynamics qualitatively resembles the dynamics shown in Section 2.2 (Fig.5).

In several cell types, such as osteoclasts, large collections of podosomes exhibit a transition to an expanding multi-podosome ring structure. The ring is densely populated with podosomes, has a width of a few podosomes and expands at a speed of ∼ 1 − 2 *µm*/*min* [44,45]. The podosomes in the multi-podosome ring are immobile and the ring’s outward expansion is achieved by a treadmilling manner: the podosomes decay at the inner part of the multi-podosome ring and form preferentially at its outer edge [43]. These multi-podosome rings move outward and merge with each other until they reach the cell periphery, where they may stabilize as a podosome belt (“sealing zone”) [46]. In certain cells the rings seem to originate at the same locations at roughly regular intervals [45].

Within the current model we can not account for the detailed dynamics of the podosomes within the expanding ring, but we can speculate that its outwards expansion may be related to the mechanism that drives the expansion of the actin ring described in Section 2.1: When a podosome in the ring decays, its constituent proteins diffuse away, and due to the ring-link deformation of the membrane the curved proteins tend to aggregate more strongly at the outer edge of the ring compared to its inner edge (see Fig.1). This mechanism will therefore naturally increase the rate of podosome formatino on the outer edge of the ring, compared to its inner edge, and over time cause the outwards expansion of the podosome ring. This expansion will depend on the rate of podosome initiation and decay, that enables this effective “podosome treadmill”.

## 3. Discussion

The results of our model can give a very natural explanation to several puzzling features observed in experiments. One such feature is that actin waves at the cell-substrate interface are observed to expand as a single ring of actin polymerization, but beyond a certain size an inner ring of actin that follows the outer one appears. The inner ring is often weaker than the outer ring, as our model predicts. This feature was first noted by Vicker [24] in *Dictyostelium discoideum* amoebae, and more recently this feature was studied in great detail [8,25].

The actin fronts observed in these experiments are very often broken and fragmented, with numerous breaks appearing along the ring [24,25]. This feature is a natural consequence in our model of the sensitivity of the actin concentration to the surface topography (Figs.3,4), due to the curvature sensitivity of the actin nucleators. Our model further predicts that the actin polymerization will tend to concentrate where the substrate has concave corners, as along the sides of elevated ridges (Fig.4). This prediction is in agreement with observations of crawling *Dicty* on surfaces with patterned ridges [47].

To conclude, we have shown some aspects of the dynamics of the actin-membrane system, when driven by convex actin activators and confined by the substrate. We show that the confinement itself provides a source of negative feedback that can drive propagating fronts. These simulations are simplified and contain several assumptions, which may not apply to all the biological cases. Furthermore, we do not claim that the actin waves along the basal membranes of cells are driven solely by the mechanism that we describe. Clearly complex reaction-diffusion feedbacks play an important role in the propagation of actin waves in cells [3,8,24,25,48,49,50]. Our work may motivate further studies of models that include both the reaction-diffusion dynamics and the membrane shape, coupled by curved membrane complexes that nucleate actin polymerization [14]. Since reaction diffusion models lead to rich dynamics, as does the curvature-actin coupling [18], we expect that models with both features could open up new classes of cell membrane dynamics to explore.

Regarding localized structures at the basal membrane, our model predicts that these may be stabilized by the formation of adhesion around the actin-core, as observed in podosomes. Furthermore, if the membrane can be maintained in a curved shape at the tip of the protrusion (rather than flatten), we predict that the curved activators will be less strongly dispersed (if at all), and the lifetime of the localized protrusion extended. This prediction is in agreement with observations that podosomes preferentially form along grooves where the membrane naturally has the curvature at the protrusion tip [51,52]. When the protrusion is able to penetrate into the substrate, as occurs in invadopodia [53], the curvature at the protrusion tip is also maintained and this can stabilize the protrusion.

## 4. Materials and Methods

### 4.1. Model without membrane-substrate adhesion

The model has two variables, 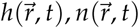, that describe the local height deformation of the membrane from its uniform state, and the local density of the membrane-bound, and curved, activators of actin polymerization, respectively. We will work in the limit of small membrane deformations, which allow us to treat the elastic energy of the membrane due to tension and bending in the quadratic limit [15,16]. This is an approximation which may be justified due to the presence of the confining boundary that naturally limited the amplitude of the membrane deformations. Non-linear effects arise in our calculations only from the conservation of the activator field *n*. For simplicity we do not consider the process of binding and unbinding of the actin activators from the membrane, which can be added in the future [10].

We start with the free energy of the membrane and actin activators [15,16]

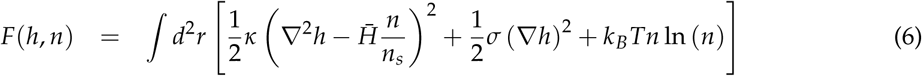

where *σ* is the effective membrane tension, *κ* the bending modulus, 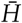 the spontaneous curvature of the actin activators and *n*_*s*_ their saturation density. In this treatment the two dimensional Laplacian ∇^2^*h* is the local mean membrane curvature, and we keep the entropic term only at the lowest order, valid for low protein densities *n* ≪ *n*_*s*_.

We model the external barrier (substrate) as a one sided harmonic potential (Fig.8) that affects the membrane if its height coordinate *h* exceeds the barrier height coordinate *h*_wall_. We also subject the membrane to a similar force (though smaller in magnitude) if its height is lower than the initial height (at *h* = 0) to account for the overall average rigidity of the cortical cytoskeleton. The wall and cytoskeleton interactions can be inserted as potentials to the free energy of the system, in the form

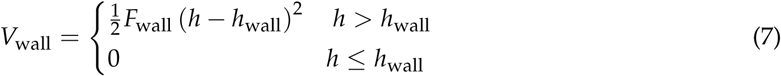

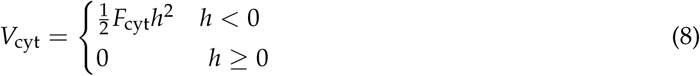

where *F*_wall_,*F*_cyt_ determine the stiffness of these potentials.

The equation of motion for the membrane height is given by [15,54]

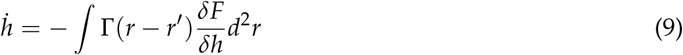

where Γ(*r* − *r*′) is the Oseen tensor for the hydrodynamic interactions through the surrounding fluid. We replace this long-range interaction kernel with a local on, as is often used for membranes that are highly confined by the cytoskeleton (and here also by the substrate) [15,16]. We therefore take: Γ(*r* − *r*′) = *µδ*(*r* − *r*′), where *µ* is the drag coefficient of the membrane. We can now use Eqs.6,9 to write the equation of motion for the membrane height

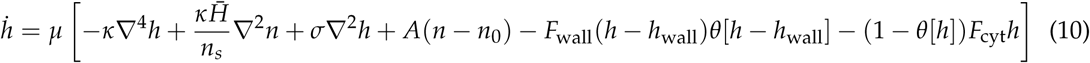

where we used the step function *θ*[*h*] to implement the truncated forces of the confining potentials of Eqs.7 and 8. The fourth term on the r.h.s. denotes the force due to actin polymerization, which is proportional to the density of actin activators with proportionality factor *A*. We subtract the average density of the actin cortex in this term to denote the fact that the cell membrane tends to be pushed away from the substrate by a layer of extracellular molecules (called the glycocalix) [55,56]. These molecules act as molecular cushions and maintain weak (non-specific) cell-substrate adhesion, and maintain osmotic pressure on the cell membrane.

The dynamics of the actin activators is given by [15,16]

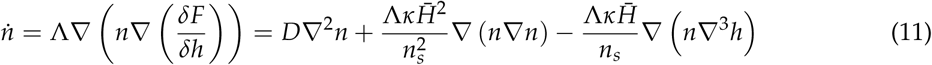

where Λ is the mobility of the activators in the membrane, and *D* = Λ*k*_*B*_*T* is their diffusion coefficient.

We note that the model presented here considers activators that are permanently bound to the membrane and respond to the curvature by flowing in the membrane. Alternatively, the activators can be considered to adsorb to the membrane from the cytoplasm in a curvature-dependent manner [10].

Linear stability analysis of these equations of motion (Eqs.10,11) indicate that the system is unstable to small perturbations in either the membrane shape or activators density, for a range of parameters. For negligible membrane tension the most unstable wavelength is

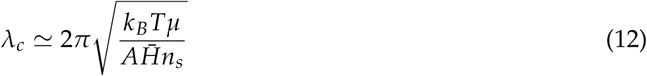

We chose the parameters to have this wavelength of order 1 *µ*m.

The numerical simulations were done using an explicit finite difference scheme centered in space and forward in time, with periodic boundary conditions. The cartesian grid used in the simulations was 0.025 × 0.025*µ*m in size.

### 4.2. Model with membrane-substrate adhesion

When including adhesion proteins, the free energy of the system becomes

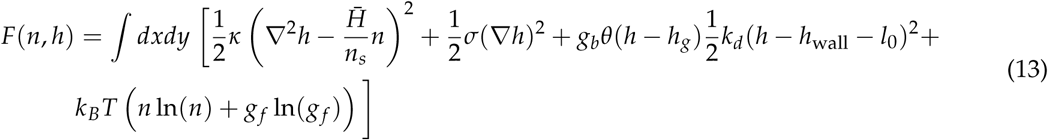

where where *l*_0_ is the length of the external part of the adhesion protein and *h*_*g*_ is the minimal membrane height required for adhesion to take place. We take this value to be *h*_*g*_ ≃ *h*_wall_ − 2*l*_0_ to allow for variations in the proteins length or membrane fluctuations.

In addition we assume that the mobility of the actin activators to decrease in regions that have high concentration of adhered adhesion proteins. The reasoning behind this is that the scaffold of plaque proteins surrounds the actin core and restricts the movement of actin filaments, and forms a “diffusion-barrier”. If actin activators are attached to actin filaments then they too will be restricted. We therefore take the mobility to decrease with the local density of bound adhesion proteins, as follows

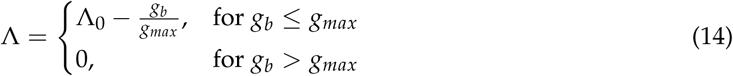

Under these conditions we can write the following equations of motion, including the dynamics of the free an bound adhesion proteins (*g*_*f*_, *g*_*b*_ respectively)

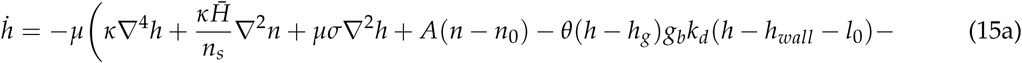

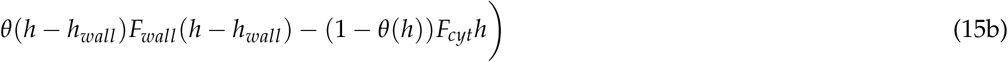

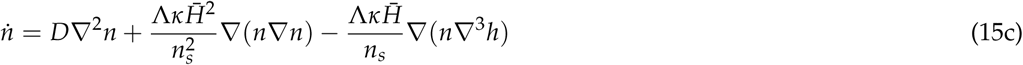

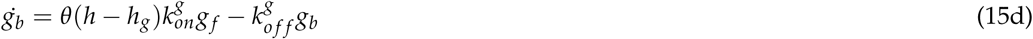

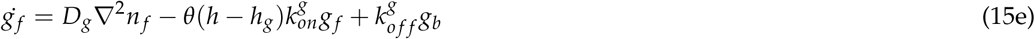

Where *D*_*g*_ is the diffusion coefficient of the free (un-adhered) adhesion proteins in the membrane, and the binding/unbinding rates of the adhesion proteins is given in Eqs.4,5.

## Author Contributions

Conceptualization, M.N. and N.S.G.; numerical simulations, M.N.; writing, M.N. and N.S.G.

## Funding

This research was funded by NAME OF FUNDER grant number XXX.” https://search.crossref.org/funding, any errors may affect your future funding.

## Acknowledgments

N.S.G is the incumbent of the Lee and William Abramowitz Professorial Chair of Biophysics. This work is made possible through the historic generosity of the Perlman family.

## Conflicts of Interest

The authors declare no conflict of interest. The funders had no role in the design of the study; in the collection, analyses, or interpretation of data; in the writing of the manuscript, or in the decision to publish the results.

